# Aging-Associated Decline in Macrophage STAT6-OXPHOS Programs Promotes Tumor-Like Multinucleated Syncytia

**DOI:** 10.64898/2026.05.18.726012

**Authors:** Li-Ying Wu, Hung-Chun Liao, Chien-Chin Chen, Chih-Wei Chou, Tim Hui-Ming Huang, Chia-Nung Hung

## Abstract

Aging can alter macrophage functions through changes in intracellular processing, mitochondrial activity, and chronic inflammatory activation; however, whether aging-associated macrophage deregulation contributes to tumor-associated multinucleated syncytial formation remains poorly understood. Here, we investigated the role of aging macrophages in promoting tumor-like multinucleated syncytia and explored the underlying metabolic mechanisms. Immunohistochemical analyses of metastatic tissue sections from patients with prostate, breast, and lung cancers demonstrated enrichment of CD68+/panCK+ multinucleated tumor-like osteoclast syncytia in elderly patients. Using *ex vivo* co-culture systems, aged bone marrow-derived macrophages exhibited significantly increased propensity to generate multinucleated syncytia containing proliferative Ki67-positive cancer-associated nuclei. These syncytia displayed attenuated mitochondrial oxidative phosphorylation (OXPHOS) programs characterized by reduced oxygen consumption rates and decreased expression of mitochondrial respiratory proteins, such as ATP5a and SDHB. Pharmacologic inhibition of STAT6 further enhanced syncytial formation and suppressed OXPHOS-associated programs, whereas treatment with the EP2 antagonist C52 partially restored mitochondrial gene expression and reduced syncytial formation. Together, these findings identify a previously unrecognized aging-associated mechanism linking macrophage deregulation, attenuated STAT6-associated mitochondrial programs, and tumor-like multinucleated syncytial formation.

## 1 Introduction

Macrophages are highly plastic innate immune cells that play a central role in tissue homeostasis, phagocytic clearance, wound repair, and inflammatory regulation (Mosser, Hamidzadeh, & Goncalves, 2021; Wculek, Dunphy, Heras-Murillo, Mastrangelo, & Sancho, 2022; Wynn & Vannella, 2016). In addition to their canonical immune functions, macrophages also actively participate in tissue remodeling through interactions with stromal, epithelial, and malignant cells in microenvironments (de Visser & Joyce, 2023; Kloosterman & Akkari, 2023). Increasing evidence suggests that aging profoundly alters macrophage biology, characterized by chronic inflammation, incomplete phagocytic processing, mitochondrial deficiency, and attenuated intracellular degradation programs (Moss, Phipps, Wilson, & Kiss-Toth, 2023; Oishi & Manabe, 2016; Seegren et al., 2023). In contrast to aging-related changes observed in circulating monocytes, tissue-associated macrophages appear particularly susceptible to age-dependent metabolic and lysosomal dysfunction because of their prolonged exposure to inflammatory and oxidative microenvironments (Lopez-Otin, Blasco, Partridge, Serrano, & Kroemer, 2023; Oishi & Manabe, 2016; M. D. Park, Silvin, Ginhoux, & Merad, 2022; Seegren et al., 2023). These aging-associated alterations have been implicated in diverse pathological conditions, including cancer progression (Bied, Ho, Ginhoux, & Bleriot, 2023; de Visser & Joyce, 2023; Fane & Weeraratna, 2020). Nevertheless, the contribution of aging macrophages to tumor cell plasticity and multinucleated syncytial formation remains to be clarified.

Recent studies, including our own, have identified macrophage-tumor hybrid cells as a previously underappreciated population capable of acquiring both immune and tumor-associated properties (Chou et al., 2023; Gast et al., 2018; Zucker et al., 2026). Several groups have reported that macrophage-tumor fusion events may contribute to metastatic dissemination, immune evasion, and enhanced cellular plasticity in multiple cancer types (Chou et al., 2023; Cozzo, Coleman, & Hursting, 2023; Tanjak et al., 2024; Zucker et al., 2026). In our previous study, we demonstrated that tumor cells undergoing macrophage-mediated phagocytic engulfment can evade complete lysosomal degradation and subsequently initiate fusion-like programs, resulting in viable hybrid cells with enhanced metastatic phenotypes (Chou et al., 2023). This process was initiated through incomplete phagocytosis, in which engulfed tumor cells retained partial viability and progressively acquired macrophage-associated characteristics (Chou et al., 2023). These observations suggested that phagocytosis is not always a terminal event for tumor cells; under certain conditions, it may instead generate viable multinucleated syncytial states capable of facilitating metastatic adaptation.

We hypothesized that aging may further potentiate this phenomenon. Senescent or stressed tumor cells frequently exhibit altered membrane signaling and impaired “don’t eat me” pathways, rendering them susceptible to macrophage engulfment (Logtenberg, Scheeren, & Schumacher, 2020; Schloesser et al., 2023). Under physiologic conditions, activated macrophages efficiently digest internalized cellular material through tightly regulated lysosomal and metabolic programs (Wculek et al., 2022; Wong et al., 2017). However, aging macrophages undergo progressive functional remodeling and exhibit reduced intracellular processing capacity (Lopez-Otin et al., 2023; Van den Bossche & Leenen, 2021). In particular, STAT6 signaling, which regulates macrophage homeostasis, anti-senescent programs, and mitochondrial fitness may become attenuated during aging (Zhou et al., 2026; Zhou et al., 2024). We speculated that reduced pSTAT6 activity in aged macrophages compromises efficient tumor-cell digestion, thereby promoting persistence of incompletely engulfed tumor cells and enhancing macrophage-tumor fusion events. Because osteoclasts arise through homotypic fusion of monocyte/macrophage lineage cells into multinucleated giant syncytia (Ma et al., 2021; Y. Park, Sato, & Lee, 2023; Sabe, Yahara, & Ishii, 2024), we further hypothesized that aging-associated macrophage remodeling may facilitate generation of tumor-like osteoclast syncytia (TOS) with enhanced proliferative and osteolytic properties in the bone microenvironment.

In this study, we demonstrate that CD68+/panCK+ tumor-like osteoclast syncytia (TOS) are enriched in bone metastatic lesions from elderly patients. Using macrophage-tumor co-culture systems, we further show that aged macrophages exhibit increased propensity to generate multinucleated syncytia characterized by attenuated mitochondrial oxidative phosphorylation (OXPHOS) programs and altered STAT6-associated signaling. These syncytial structures retained proliferative tumor-associated nuclei, supporting acquisition of tumor-like cellular properties following macrophage-mediated engulfment and fusion-related processing events. Taken together, our findings identify a previously unrecognized aging-associated mechanism linking macrophage remodeling, attenuated intracellular processing capacity, and formation of tumor-like multinucleated syncytia.

## 2 Results

### 2.1 Aging-Associated Enrichment of TOS in Human Bone Metastatic Lesions

Immunohistochemical (IHC) analysis was performed using the macrophage marker CD68 and the epithelial tumor marker pan-cytokeratin (panCK) to identify putative TOS at tumor-bone interface regions in bone metastatic tissue sections from 32 patients with breast, prostate, and lung cancers (Table S1). The workflow and scoring methodology are illustrated in Figure 1A: 20-100 microscopic fields per patient specimen were analyzed to quantify CD68+/panCK+ multinucleated cells, which were stratified into four semi-quantitative categories (0-III) according to the abundance of bone-resorbing syncytia. An integrated bone resorbing cell index was subsequently calculated using weighted category scores. Representative IHC images demonstrated CD68+/panCK+ syncytia localized predominantly at tumor-bone interface regions in metastatic lesions from prostate, breast, and lung cancers (Figure 1B, *upper panels*). These dual-positive cells frequently displayed multinucleated morphology and were enriched in areas adjacent to bone trabeculae, supporting their osteoclast-like phenotype. In several specimens, these multinucleated structures exhibited irregular giant-cell morphology closely resembling activated osteoclasts within osteolytic lesions (Figure 1B, *lower panels*). Higher-magnification images further demonstrated that these structures exhibited an irregular, multinucleated morphology with strong co-expression of CD68/panCK, supporting their identity as tumor-associated osteoclast-like syncytia (Figure 1C). Notably, specimens from older patients (≥65 years old) exhibited substantially higher proportions of Category II-III lesions than younger patients (<65 years old), who more frequently showed Category 0-I lesions (Figure 1D). Consistently, the bone-resorbing cell index was significantly elevated in metastatic tissues from older patients compared with younger patients (*p* < 0.01; Figure 1E). Collectively, these findings suggest that aging is associated with increased accumulation of tumor-like syncytia within metastatic bone lesions across multiple epithelial cancer types.

**FIGURE 1.**
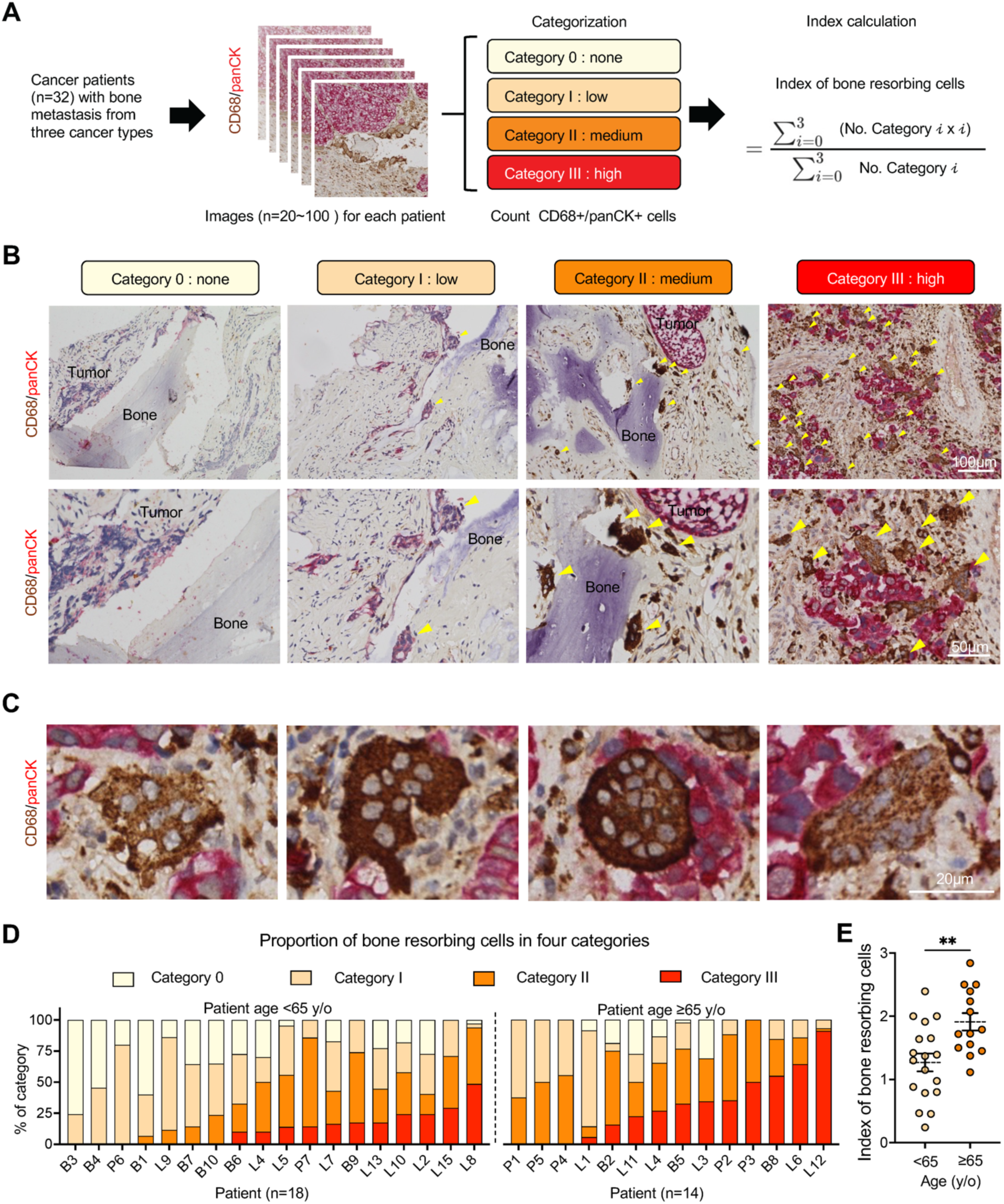
Increased accumulation of bone-resorbing cells in bone metastatic tissues from older patients. (A) Schematic illustration of the workflow used to evaluate bone-resorbing cells in metastatic bone tissue samples obtained from 32 patients with bone metastases derived from three cancer types. CD68/panCK double immunohistochemical staining was performed, and 20-100 images were analyzed for each patient. CD68+/panCK+ cells located adjacent to bone regions were quantified and classified into four categories based on cell abundance: Category 0 (none, zero bone-resorbing cells), Category I (low, 1-4 bone-resorbing cells), Category II (medium, 5-9 bone-resorbing cells), and Category III (high, ≥10 bone-resorbing cells). A bone-resorbing cell index was calculated using the weighted scoring formula shown. (B) Representative images of CD68/panCK double immunohistochemical staining corresponding to each classification category. Yellow arrowheads indicate CD68+/panCK+ bone-resorbing cells adjacent to bone regions. Scale bars, 100 μm (upper panel) and 50 μm (lower panels). (C) Representative high-magnification images of CD68/panCK double-positive cells in metastatic bone tissues. Scale bars, 20 μm. (D) Distribution of bone-resorbing cell categories in individual patients stratified by age (<65 years, n = 18; ≥65 years, n = 14). Each bar represents the proportion of images assigned to each category for an individual patient. (E) Quantification of the bone-resorbing cell index in patients younger than 65 years and those aged 65 years or older. Data are presented as mean ± SEM (<65 years, n = 18; ≥65 years, n = 14). Statistical significance was determined by the Mann-Whitney test. *\*\*P < 0.01*.

### 2.2 Aging-Associated Macrophages Promote the Formation of Multinucleated Syncytia

Given the increased presence of TOS in bone metastatic lesions from older patients, we next investigated whether aging-associated macrophages intrinsically promote formation of multinucleated tumor syncytia using an *ex vivo* bone marrow-derived macrophage (BMDM) co-culture system (Figure 2A). Bone marrow cells isolated from young (5-week-old) or aged (86-week-old) mice were differentiated into macrophages prior to co-culture with fluorescently labeled TRAMP-C2 cancer cells. Flow cytometric gating strategies confirmed efficient macrophage maturation, with approximately 80% of viable cells exhibiting a CD11b+F4/80+ macrophage phenotype in both age groups (Figure S1A, B).

**FIGURE 2.**
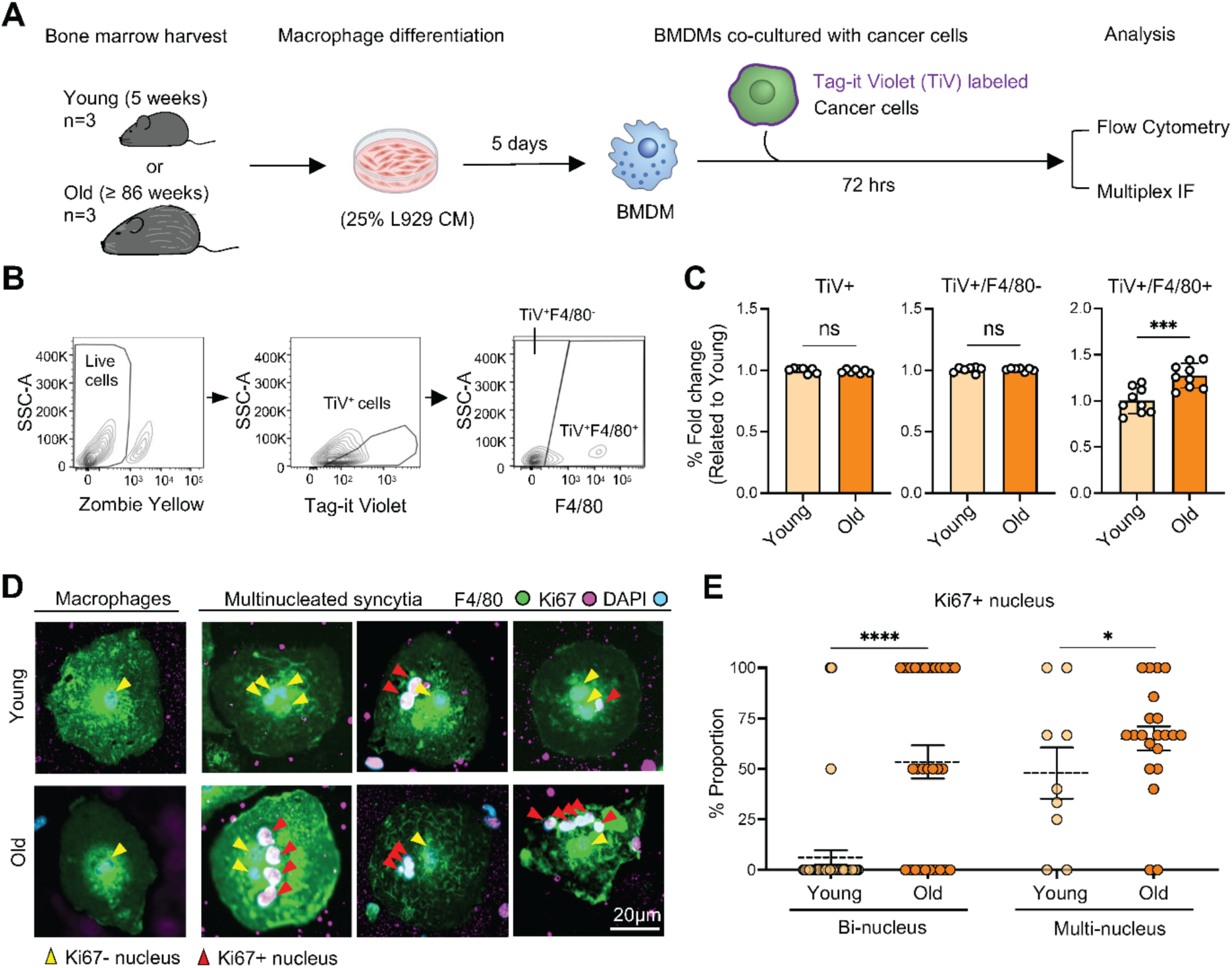
Age-related enhancement of macrophage-tumor cell fusion and multinucleated syncytia formation. (A) Schematic illustration of the experimental workflow. Bone marrow cells isolated from young (5-week-old) and aged (≥86-week-old) mice (n = 3 per group) were differentiated into bone marrow-derived macrophages (BMDMs) in medium containing 25% L929 conditioned medium for 5 days. BMDMs were subsequently co-cultured with Tag-it Violet (TiV)-labeled cancer cells for 72 h prior to flow cytometric and multiplex IF analyses. (B) Representative flow cytometry gating strategy used to identify live cells, TiV+ cells, TiV+/F4/80-tumor cells, and TiV+/F4/80+ fused cells. (C) Quantification of TiV+ cells, TiV+/F4/80-cells, and TiV+/F4/80+ fused cells following co-culture of young or aged BMDMs with cancer cells. Data are presented as fold change relative to the young group (n = 9 per group). (D) Representative immunofluorescence images of macrophages and multinucleated syncytia stained for F4/80, Ki67, and DAPI following co-culture with young or aged BMDMs. Yellow arrowheads indicate Ki67-negative nuclei, and red arrowheads indicate Ki67-positive nuclei within multinucleated syncytia. Scale bar, 20 μm. (E) Quantification of the proportion of bi-nucleated (n = 41 in young group and n = 29 in old group) and multinucleated syncytia (n = 9 in young group and n = 21 in old group) containing Ki67-positive nuclei in co-cultures with young or aged BMDMs. Data are presented as mean ± SEM. Statistical significance was determined by the Mann-Whitney test. *ns, not significant; *P < 0.05; ***P < 0.001*.

Using Tag-it Violet (TiV)-labeled tumor cells, flow cytometric analyses demonstrated significantly increased TiV+/F4/80+ cell populations in co-cultures containing aged BMDMs compared with young BMDMs (*p* < 0.001, Figure 2B, C). In contrast, TiV+/F4/80-tumor cell fractions showed less pronounced age-dependent differences, suggesting that aging preferentially enhances macrophage-tumor syncytial formation rather than simply promoting tumor-cell survival.

Representative multiplex immunofluorescence (IF) analyses further revealed abundant multinucleated F4/80+/Ki67+ syncytia within aged co-cultures (Figure 2D, *upper panel*). Importantly, Ki67 positivity within OLS supports a fusion-related origin involving proliferative tumor-derived nuclei, as parental BMDMs alone did not express detectable Ki67 signals (Figure 2D, *lower panel*). Morphologically, these multinucleated structures closely resembled the CD68+/panCK+ TOS identified in human bone metastatic tissues.

We next quantified the proliferative characteristics and multinucleation status of co-cultured cells derived from young or aged BMDMs. Specifically, the proportion of Ki67+ nuclei was analyzed in bi-nucleated and multi-nucleated syncytia. Aged BMDM-derived syncytia exhibited significantly increased proportions of

Ki67+ nuclei compared with syncytia generated from young macrophages in both bi-nucleated and multi-nucleated populations (*p* < 0.001 and *p* < 0.05, respectively; Figure 2D, E). These findings suggest that multinucleated syncytia formed under aging conditions retain enhanced proliferative activity despite acquiring multinucleated osteoclast-like morphology. This increase was particularly prominent in multi-nucleated syncytia, implicating that aging-associated macrophages not only promote syncytial formation but also support maintenance of proliferative cancer-associated nuclei within enlarged bone-resorbing structures. Collectively, these observations support our clinical observations in which aging macrophages facilitate generation of proliferative TOS in the bone microenvironment.

### 2.3 STAT6 Inhibition Enhanced the Formation of Multinucleated Syncytia

Because aging-associated macrophages demonstrated enhanced TOS, we next investigated whether impaired STAT6 signaling contributes to this process using the pharmacologic STAT6 inhibitor AS1517499 (2 μM) (Figure 3A)(Zhou et al., 2024). Young BMDMs from 5-week-old mice or RAW264.7 macrophage-like preosteoclasts (Herb et al., 2024) were co-cultured with multiple murine cancer cell lines, including TRAMP-C2, Py8119, Py230, LLC1, Myc-CaP, and mKRC.1, followed by flow cytometric and multiplex IF analyses (Figure 3B).

**FIGURE 3.**
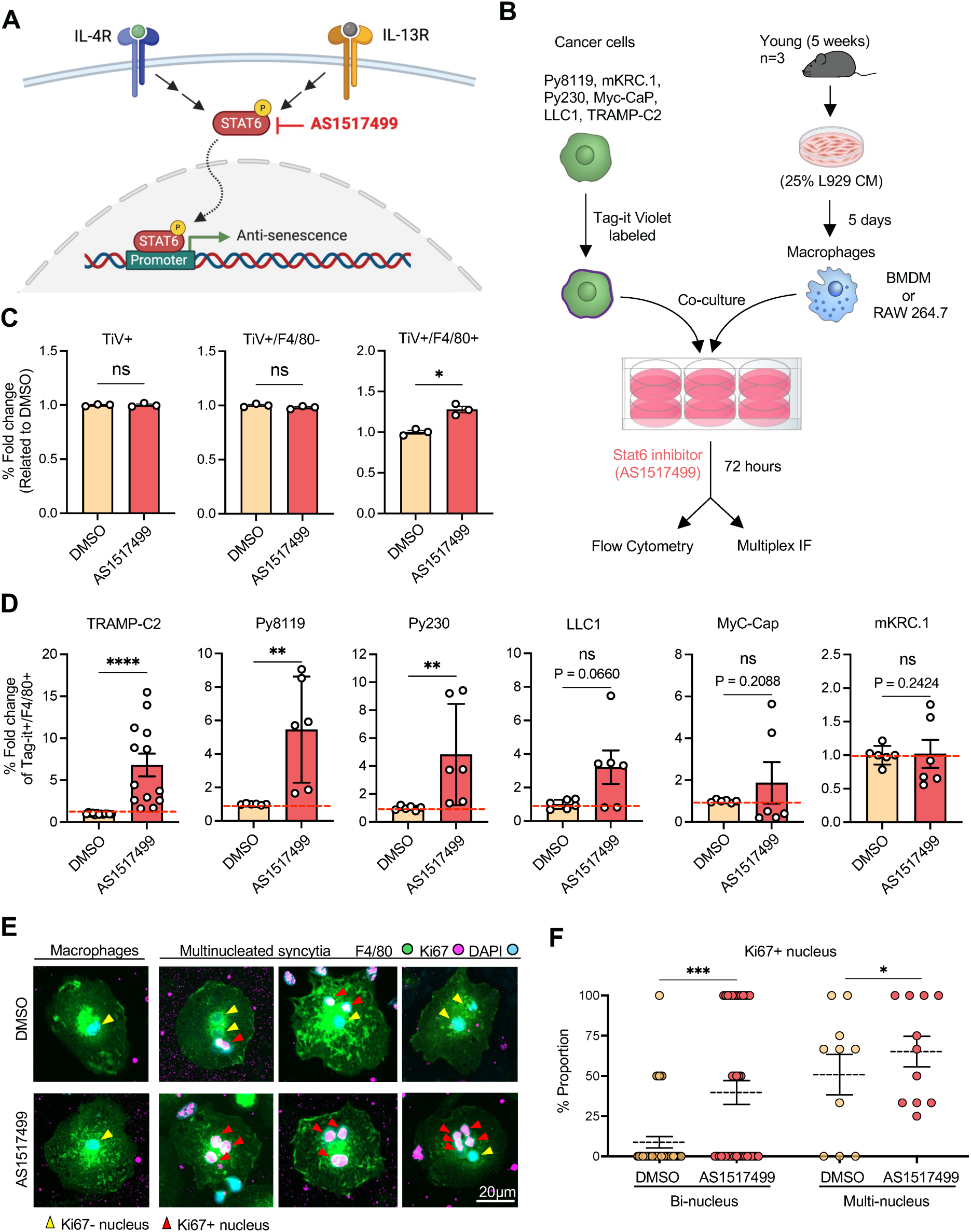
STAT6 inhibition enhances macrophage-tumor cell fusion and multinucleated syncytia formation. (A) Schematic illustration of STAT6 inhibition by AS1517499. IL-4R and IL-13R signaling activate STAT6-mediated transcriptional regulation, which was blocked using the STAT6 inhibitor AS1517499. (B) Experimental workflow for macrophage-tumor cell fusion assays. Tag-it Violet-labeled cancer cells (Py8119, mKRC.1, Py230, MyC-CaP, LLC1, and TRAMP-C2) were co-cultured with BMDMs or RAW264.7 macrophages in the presence of DMSO or AS1517499 for 72 h prior to flow cytometric and multiplex immunofluorescence analyses. (C) Flow cytometric quantification of TiV+ cells, TiV+/F4/80-cells, and TiV+/F4/80+ fused cells after treatment with DMSO or AS1517499. Data are presented as fold change relative to the DMSO control (n = 3 per group). (D) Quantification of TiV+/F4/80+ fused cells generated by co-culture of macrophages with different cancer cell lines (TRAMP-C2, Py8119, Py230, LLC1, MyC-CaP, and mKRC.1) in the presence of DMSO or AS1517499. Data are presented as fold change relative to the DMSO control (n ≥ 6 per group). (E) Representative immunofluorescence images of macrophages and multinucleated syncytia stained for F4/80, Ki67, and DAPI following DMSO or AS1517499 treatment. Yellow arrowheads indicate macrophage nuclei, and red arrowheads indicate Ki67-positive nuclei. Scale bar, 20 μm. (F) Quantification of the proportion of bi-nucleated (n = 40 in DMSO group and n = 39 in AS1517499 group) and multinucleated syncytia (n = 10 in DMSO group and n = 11 in AS1517499 group) containing Ki67-positive nuclei in DMSO- or AS1517499-treated co-cultures. Data are presented as mean ± SEM. Statistical significance was determined by the Mann-Whitney test. *ns, not significant; *P < 0.05; **P < 0.01; ***P < 0.001; ****P < 0.0001*.

Flow cytometric analyses demonstrated a trend toward increased formation of TiV+/F4/80+ syncytial populations following STAT6 inhibition. Although total TiV+ tumor-cell populations and TiV+/F4/80-fractions showed minimal changes following AS1517499 treatment, TiV+/F4/80+ populations were significantly increased after STAT6 inhibition (*p* < 0.05, Figure 3C). These findings suggest that reduced STAT6 referentially can promote macrophage-tumor syncytial formation rather than altering general tumor-cell viability. Across multiple tumor models, including TRAMP-C2, Py8119, and Py230, AS1517499 treatment significantly increased generation of TiV+/F4/80+ populations (*p* < 0.001, *p* < 0.01, and *p* < 0.01, respectively; Figure 3D), whereas more modest and non-significant effects were observed in LLC1, Myc-CaP, and mKRC.1 co-cultures. These findings further suggest that susceptibility to syncytial formation may vary according to tumor lineage or intrinsic tumor-cell properties.

Representative multiplex IF analyses further demonstrated that AS1517499-treated cultures exhibited a visibly increased number of multinucleated syncytia containing abundant Ki67+ tumor-associated nuclei relative to DMSO-treated controls (Figure 3E). Quantification of Ki67+ nuclei further demonstrated significant increases in both bi-nucleated and multi-nucleated populations following STAT6 inhibition (*p* < 0.001 and *p* < 0.05, respectively; Figure 3F). Because STAT6 signaling has been implicated in macrophage homeostasis, anti-senescent regulation, and intracellular processing functions(Zhou et al., 2026; Zhou et al., 2024), these results support the hypothesis that aging-associated attenuation of STAT6 activity may reduce efficient degradation of engulfed tumor cells, thereby facilitating persistence of incompletely processed tumor material and promoting syncytial fusion events.

### 2.4 Aging-Associated Multinucleated Syncytia Exhibit Reduced Mitochondrial OXPHOS Programs

Because attenuation of STAT6 signaling enhanced multinucleated syncytia formation in co-cultures, we next investigated whether aging-associated macrophage fusion events are accompanied by alterations in mitochondrial oxidative phosphorylation (OXPHOS) programs (Figure 4A) (Minhas et al., 2019; Seegren et al., 2023; Van den Bossche & Leenen, 2021). Given that efficient lysosomal digestion and intracellular processing require substantial mitochondrial activity, we determined whether aging macrophages exhibit diminished metabolic fitness that favors persistence of incompletely processed tumor material and promotes syncytial formation. Quantitative RT-PCR analyses demonstrated that aged BMDMs exhibited reduced expression of several OXPHOS-associated genes compared with young macrophages, including *Gpx1* and *Cox5a*, whereas *Sdhb*, *Cox2*, *Atp5a*, and *Atp5d* showed more modest or non-significant reductions (Figure 4B). These findings suggest partial suppression of mitochondrial respiratory programs during macrophage aging.

**FIGURE 4.**
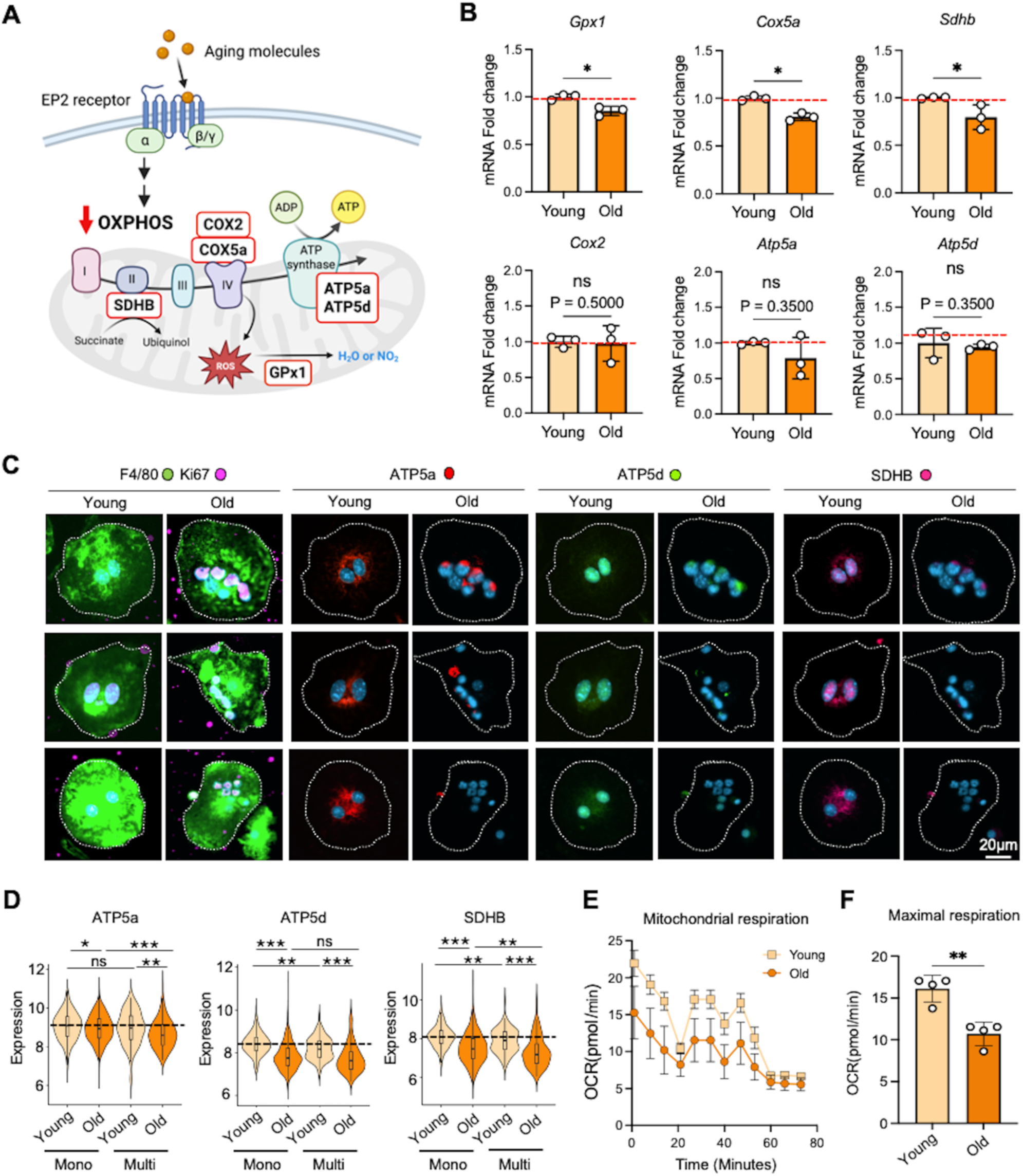
OXPHOS-associated mitochondrial function is reduced in macrophage-tumor fusion cells from aged mice. (A) Schematic illustration of aging-associated suppression of OXPHOS signaling in macrophages. Representative OXPHOS-related proteins analyzed in this study, including COX2, COX5a, SDHB, ATP5a, ATP5d, and GPX1, are indicated within the mitochondrial respiratory chain. (B) Quantitative PCR analysis of OXPHOS-associated genes, including *Gpx1, Cox5a, Sdhb, Cox2, Atp5a,* and *Atp5d*, in BMDMs isolated from young and aged mice. Gene expression levels were normalized to the young group. Data are presented as mean ± SEM (n = 3 per group). (C) Representative multiplex immunofluorescence images of macrophage-tumor fusion cells generated from young or aged mice stained for F4/80, ATP5a, ATP5d, and SDHB. Dashed lines indicate cell boundaries. Scale bar, 20 μm. (D) Quantification of ATP5a, ATP5d, and SDHB expression in mononucleated (Mono) and multinucleated (Multi) macrophage-tumor fusion cells derived from young and aged mice. Data are presented as violin plots showing distribution and median values. (E) Seahorse extracellular flux analysis showing mitochondrial respiration profiles of macrophage-tumor fusion cells derived from young and aged mice (n =3 per group). Oxygen consumption rate (OCR) was monitored over time following sequential treatment with mitochondrial stress test reagents. (F) Quantification of maximal respiration capacity in macrophage-tumor fusion cells from young and aged mice based on Seahorse analysis. Data are presented as mean ± SEM. Statistical significance was determined by the Mann-Whitney test. \**P* < 0.05; \*\**P* < 0.01; \*\*\**P < 0.001; ns, not significant*.

To further examine mitochondrial status in multinucleated syncytia, multiplex IF analyses were performed on syncytia generated from young and aged macrophage co-cultures. Representative images showed reduced expression of mitochondrial respiratory proteins ATP5a, ATP5d, and SDHB in multinucleated syncytia derived from aged macrophages compared with those generated from young macrophages (Figure 4C). Quantitative analyses further demonstrated significant reductions of ATP5a and SDHB expression in both mononucleated and multinucleated syncytial populations derived from aged macrophages, whereas ATP5d expression showed more variable changes (Figure 4D). Importantly, the reduction in mitochondrial protein expression was particularly evident in multinucleated syncytia, supporting the notion that aging-associated syncytia exhibit attenuated mitochondrial metabolic programs. Consistent with these observations, Seahorse extracellular flux analyses demonstrated markedly reduced oxygen consumption rates (OCR) and diminished maximal respiratory capacity in multinucleated syncytia generated from aged macrophages compared with young controls (*p* < 0.01, Figure 4E, F).

Because reduced STAT6 signaling promoted syncytial formation in the aforementioned co-cultures, we next examined whether STAT6 inhibition similarly altered mitochondrial programs. AS1517499 treatment of young BMDMs reduced expression of multiple OXPHOS-associated genes, including *Sdhb*, *Cox2*, *Cox5a*, and *Atp5a* (Figure S2A). In parallel, multiplex IF analyses demonstrated reduced ATP5a, ATP5d, and SDHB protein expression in multinucleated syncytia following AS1517499 treatment in co-cultures (Figure S2B, C). Collectively, these findings suggest that reduced STAT6-associated mitochondrial programs in aging macrophages may facilitate formation of metabolically dysfunctional multinucleated syncytia.

### 2.5 EP2 Antagonism Suppresses Multinucleated Syncytia Formation and Restores Mitochondrial OXPHOS Programs

Because aging-associated multinucleated syncytia exhibited attenuated mitochondrial OXPHOS programs together with enhanced fusion activity, we next investigated whether pharmacologic restoration of mitochondrial-associated pathways could suppress syncytial formation using the EP2 antagonist C52 (Figure 5A). Previous studies have implicated EP2 signaling in aging-associated inflammatory and metabolic dysfunction (Chen et al., 2022; Minhas et al., 2021); therefore, we hypothesized that EP2 inhibition may partially restore macrophage metabolic fitness and limit the formation of multinucleated syncytia in co-cultures.

**FIGURE 5.**
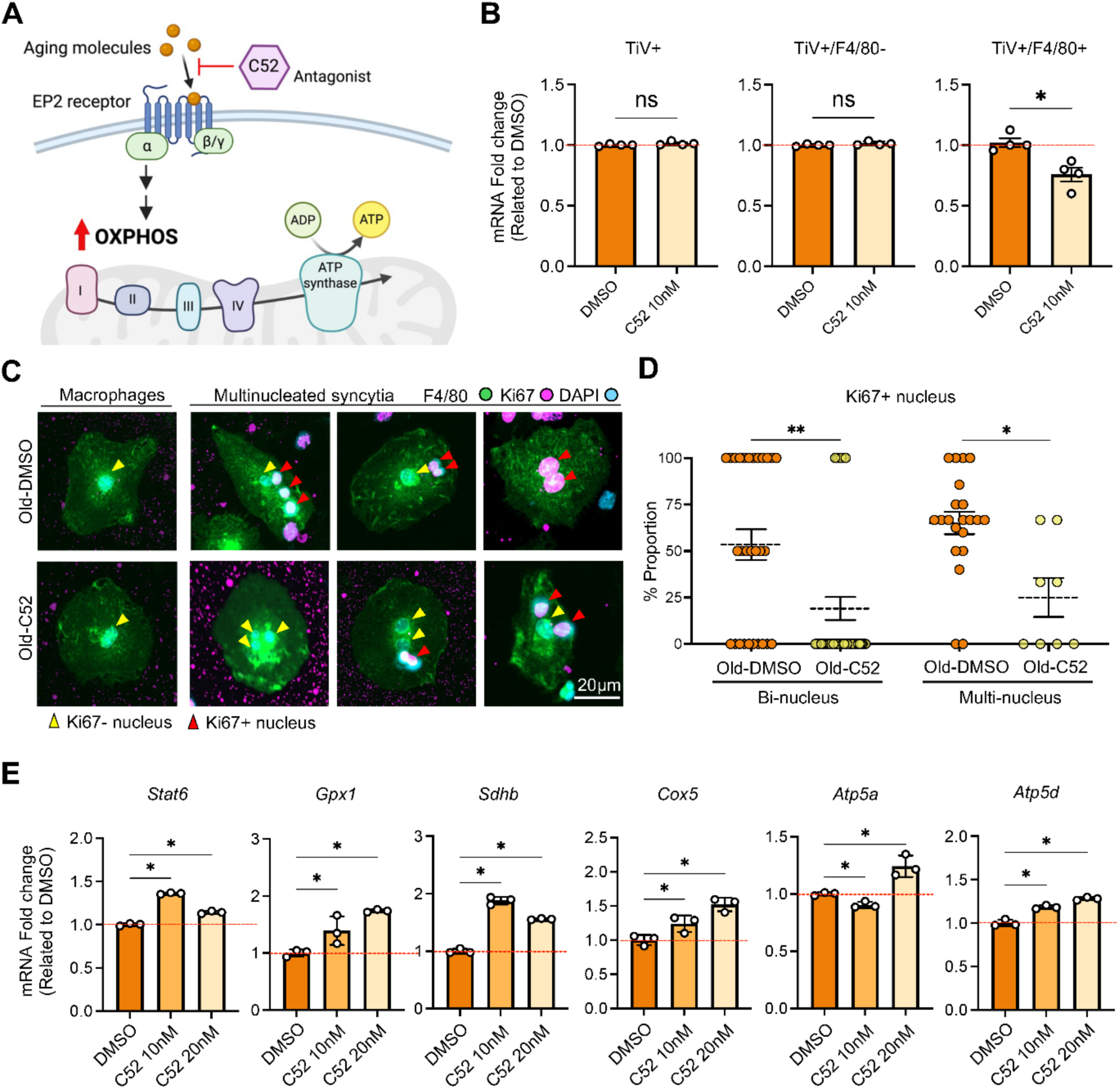
EP2 antagonist treatment modulates macrophage-tumor fusion and restores OXPHOS-associated gene expression in aged BMDMs. (A) Schematic illustration of EP2 receptor antagonism by C52. Aging-associated signaling through the EP2 receptor suppresses oxidative phosphorylation (OXPHOS)-related pathways, which were targeted using the EP2 antagonist C52. (B) Flow cytometric quantification of TiV+ cells, TiV+/F4/80- tumor cells, and TiV+/F4/80+ fused cells following co-culture of aged BMDMs with tumor cells in the presence of DMSO or C52 (10 nM). Data are presented as fold change relative to the DMSO control (n = 4 per group). (C) Representative immunofluorescence images of macrophages and multinucleated syncytia generated from aged BMDMs treated with DMSO or C52. Cells were stained for F4/80, Ki67, and DAPI. Yellow arrowheads indicate Ki67-negative nuclei, and red arrowheads indicate Ki67-positive nuclei within multinucleated syncytia. Scale bar, 20 μm. (D) Quantification of the proportion of bi-nucleated (n = 29 in DMSO group and n = 42 in C52 group) and multinucleated syncytia (n = 21 in DMSO group and n = 8 in C52 group) containing Ki67-positive nuclei in aged BMDM co-cultures treated with DMSO or C52. Data are presented as mean ± SEM. Statistical significance was determined by the Mann-Whitney test. *\*P < 0.05; **P < 0.01.* (E) Quantitative PCR analysis of Stat6 and OXPHOS-associated genes, including Gpx1, Sdhb, Cox5, Atp5a, and Atp5d, in aged BMDMs treated with DMSO, C52 (10 nM), or C52 (20 nM). Gene expression levels were normalized to the DMSO control. Data are presented as mean ± SEM (n = 3 per group). Statistical significance was determined by ordinary one-way ANOVA. *\*P < 0.05*.

Bone marrow cells isolated from aged mice were differentiated into BMDMs and subsequently co-cultured with Tag-it Violet-labeled tumor cells in the presence of DMSO or C52 (10-20 nM) prior to flow cytometric and multiplex immunofluorescence analyses (Figure S3A). Bright-field imaging demonstrated that C52 treatment visibly improved the growth and cellular expansion of aged BMDMs, possibly reflecting partial restoration of mitochondrial metabolic fitness and reduced aging-associated cellular stress (Figure S3B). Further flow cytometric gating analyses demonstrated a trend toward reduced TiV+/F4/80+ populations following C52 treatment (see representative flow cytometry plots shown in Figure S3C). Quantitative analyses demonstrated that treatment with C52 significantly reduced TiV+/F4/80+ populations in aged co-cultures (p < 0.01), whereas total TiV+ tumor-cell populations and TiV+/F4/80- fractions showed comparatively modest changes (Figure 5B). These findings suggest that EP2 antagonism preferentially suppresses macrophage-tumor syncytial formation rather than general tumor-cell viability.

Representative multiplex IF analyses further demonstrated that C52-treated co-cultures exhibited visibly reduced multinucleated structures and diminished Ki67+ proliferation-associated nuclei (Figure 5C). Quantification confirmed significant reductions in both bi-nucleated and multi-nucleated syncytia following C52 treatment (*p* < 0.01 and *p* < 0.05, respectively, Figure 5D).

To determine whether EP2 antagonism restored mitochondrial metabolic programs, quantitative RT-PCR analyses were performed on aged BMDMs treated with DMSO or C52. The C52 treatment increased expression of Stat6 and multiple OXPHOS-associated genes, including *Gpx1*, *Sdhb*, *Cox5a*, *Atp5a*, and *Atp5d* in aged macrophages (Figure 5E). Collectively, these findings suggest that EP2-associated inflammatory signaling contributes to aging-associated multinucleated syncytial formation through suppression of mitochondrial OXPHOS programs and reduced macrophage processing capacity.

## 3 Discussion

In this study, we identify an aging-associated mechanism that promotes the formation of multinucleated tumor-associated syncytia *in vitro* and further relates these structures to osteoclast-like multinucleated syncytia observed in human bone metastatic tissues. Although our clinical cohort was limited to a single retrospective tissue-based analysis, CD68+/panCK+ multinucleated syncytia were consistently enriched in bone metastatic lesions from elderly patients with prostate, breast, and lung cancers. These multinucleated structures localized predominantly near tumor-bone interface regions and morphologically resembled osteoclast-like giant cells, supporting a potential role in pathological bone remodeling during metastatic progression.

Recent studies of multinucleated giant cells (MGCs) have emphasized that macrophage-lineage cells can undergo cell-cell fusion to generate specialized multinucleated populations, including osteoclasts, foreign-body giant cells, and Langhans-type giant cells (Ahmadzadeh et al., 2023; Milde et al., 2015). These cells are generally viewed as products of macrophage-macrophage fusion that arise in response to persistent inflammatory, infectious, or foreign-body stimuli and acquire specialized degradative, tissue-remodeling, or granulomatous functions (Cai, Jiang, & He, 2024). In contrast, the tumor-like osteoclast syncytia described in our study appear to represent a distinct cancer-associated multinucleated state, characterized by co-expression of macrophage/osteoclast-lineage markers and tumor-associated epithelial features, enrichment at tumor-bone interfaces, and retention of proliferative cancer-associated nuclei. This phenotype differs from conventional MGCs because it may arise through macrophage-tumor interactions, incomplete tumor-cell processing, and fusion-related hybridization rather than macrophage-macrophage fusion alone. Consistent with recent reports that osteoclast-like giant cells can occur within tumors and may reflect tumor-associated macrophage biology (Cyrta et al., 2022; Sajjadi et al., 2022), our findings suggest that aging-associated macrophage remodeling may favor the formation of tumor-like osteoclast syncytia with potential roles in osteolytic remodeling and metastatic adaptation.

A major finding of this study is that aging macrophages exhibit substantially enhanced propensity to generate multinucleated syncytia following co-culture with cancer cells. Importantly, these syncytia contained Ki67-positive nuclei, supporting incorporation of proliferative cancer-derived nuclei into multinucleated structures, whereas parental BMDMs alone exhibited minimal or undetectable Ki67 expression. These observations provide indirect but important evidence supporting a fusion-related hybrid origin rather than simple macrophage aggregation. In addition, multinucleated syncytia generated from aged macrophages displayed attenuated mitochondrial oxidative phosphorylation (OXPHOS) programs characterized by reduced OCR and decreased expression of mitochondrial respiratory proteins, including ATP5a and SDHB. Despite reduced mitochondrial fitness, these multinucleated syncytia retained proliferative cancer-associated nuclei, suggesting acquisition of cellular features potentially favorable for tissue invasion and persistence within stressed microenvironments.

Another finding of this study is the identification of attenuated STAT6-associated signaling as a potential regulator of aging-related syncytial formation. Pharmacologic inhibition of STAT6 enhanced macrophage-tumor syncytial formation and increased proliferative multinucleated populations across multiple tumor models. In parallel, aging-associated multinucleated syncytia exhibited suppression of mitochondrial OXPHOS-associated programs, whereas treatment with the EP2 antagonist C52 partially restored mitochondrial gene expression and reduced syncytial formation. These findings suggest that aging macrophages may undergo defective intracellular processing programs characterized by reduced pSTAT6 activity and altered mitochondrial regulation, thereby permitting persistence of incompletely digested tumor cells and promoting fusion-like events. Conceptually, these observations are consistent with our prior report demonstrating that incomplete phagocytosis can initiate viable macrophage-tumor hybrid-cell formation (Chou et al., 2023).

From a clinical perspective, our findings raise the possibility that aging-associated macrophage remodeling may contribute to the formation of osteoclast-like multinucleated syncytia capable of promoting osteolytic progression in elderly patients. The enrichment of CD68+/panCK+ multinucleated structures in older patient specimens further supports the clinical relevance of this aging-associated phenotype (Figure 1). More broadly, these findings suggest that aging-related alterations in macrophage metabolic fitness and intracellular processing capacity may influence tumor cell plasticity and generation of aggressive multinucleated states within metastatic microenvironments.

This study should be considered primarily as a proof-of-principle investigation, and several limitations warrant consideration. First, the patient cohort was relatively modest and retrospective in nature. Larger prospective studies will be necessary to validate the clinicopathological significance of CD68+/panCK+ multinucleated syncytia in metastatic disease. Second, although our co-culture systems strongly support fusion-related syncytial formation, definitive lineage-tracing studies *in vivo* will be required to establish the precise cellular origin and fate of these multinucleated populations during metastatic progression. Third, the molecular mechanisms linking aging, reduced pSTAT6 activity, mitochondrial dysfunction, incomplete phagocytosis, and syncytial formation remain incompletely defined. Future studies integrating spatial transcriptomics, metabolic profiling, and genetically engineered aging models may further clarify how senescent immune microenvironments regulate tumor-associated multinucleated syncytia and metastatic adaptation within bone. Taken together, our findings support a model in which aging macrophages actively facilitate the generation of osteoclast-like multinucleated syncytia through altered intracellular processing and mitochondrial dysregulation.

## 4 Materials and Methods

### 4.1 Patients

All patients were enrolled under a protocol approved by the Ditmanson Medical Foundation Chia-Yi Christian Hospital. Written informed consent was obtained from all participants prior to sample collection. Tumor tissues were collected from 32 cancer patients with confirmed bone metastasis. The clinicopathological characteristics of the patients are summarized in Supplementary Table S1.

### 4.2 Cells

Firefly luciferase reporter-expressing Py230 and Py8119 cells (pSBE4-Luc) were generously provided by Dr. Lesley Ellies (University of California, San Diego). The mKRC.1 cell line was purchased from Applied Biological Materials (T8096). L-929 (CCL-1), LLC1 (CRL-1642), MyC-CaP (CRL-3255), RAW264.7 (TIB-71), and TRAMP-C2 (CRL-2731) cells were obtained from ATCC. LLC1 and Py8119 cells were maintained in DMEM/F-12 medium (Thermo Fisher Scientific, 11320033) supplemented with 10% fetal bovine serum (FBS; Sigma-Aldrich, F2442) and 1% penicillin-streptomycin (Thermo Fisher Scientific, 15140122). mKRC.1 cells were cultured in RPMI-1640 medium (Thermo Fisher Scientific, 11875119) containing 5% FBS and 1% penicillin-streptomycin. L-929, MyC-CaP, Py230, and RAW264.7 cells were maintained in DMEM (Thermo Fisher Scientific, 11965092) supplemented with 10% FBS and 1% penicillin-streptomycin, whereas TRAMP-C2 cells were cultured in RPMI-1640 supplemented with 10% FBS and 1% penicillin-streptomycin. All cell lines were maintained at 37°C in a humidified incubator containing 5% CO₂. For the generation of stable fluorescence/luciferase-labeled cell lines, LLC1, mKRC.1, MyC-CaP, and TRAMP-C2 cells were transduced with lentiviral particles carrying the pFUGW-Pol2-ffLuc2-eGFP construct (Addgene plasmid #14883) for 24 h. Lentiviral particles were produced using the 3rd Generation Packaging Mix from Applied Biological Materials according to the manufacturer’s instructions.

### 4.3 Immunohistochemical (IHC) Analysis

Formalin-fixed paraffin-embedded (FFPE) bone metastatic tissue specimens from patients with breast, prostate, and lung cancers were used for histological and immunohistochemical analyses following institutional approval and relevant ethical regulations. Paraffin-embedded bone sections were deparaffinized by sequential immersion in xylene (three times), followed by 100%, 90%, 70%, 50%, and 30% ethanol, and finally PBS, each for 10 min. For immunohistochemistry (IHC), patient tissue samples were sectioned into 4-μm slices and stained using the BOND-III automated IHC staining system (Leica Biosystems Newcastle Ltd., 22.2201) according to the manufacturer’s instructions. The primary antibodies used were anti-panCK (1:300, Sakura Finetek USA, 60-0022) and anti-CD68 (1:400, Zeta Corporation, 50-221-5864). Chromogenic detection was performed using Bond Polymer Refine Red Detection-AP Red (Leica Biosystems Newcastle Ltd., DS9390) and Bond Polymer Refine Detection-DAB (Leica Biosystems Newcastle Ltd., DS9800). Stained slides were scanned and imaged using a slide scanning microscope (Keyence, BZ-X800).

To evaluate tumor-associated osteoclast-like syncytia (TOS), 20-100 microscopic fields were analyzed from each patient specimen. CD68+/panCK+ multinucleated cells located at tumor–bone interface regions were quantified and classified into four semi-quantitative categories according to cell abundance: Category 0 (none, zero bone-resorbing cells), Category I (low, 1-4 bone-resorbing cells), Category II (medium, 5-9 bone-resorbing cells), and Category III (high, ≥10 bone-resorbing cells). A bone-resorbing cell index was calculated using the following formula:

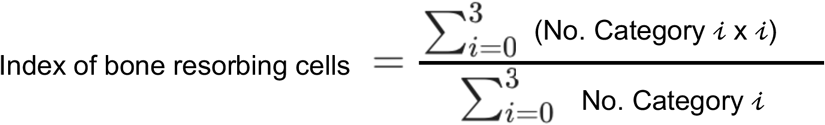

### 4.4 Bone Marrow-Derived Macrophages (BMDMs) Isolation and Differentiation

Bone marrow cells were freshly isolated from the femurs and tibias of untreated 6-8-week-old (young) and ≥86-week-old (old) C57BL/6J mice. Cells were incubated with ACK lysis buffer (Gibco) to remove red blood cells (RBCs), followed by filtration through 35 μm cell strainer cap tubes (Falcon) to obtain an RBC-free single-cell suspension. To prepare L-929 conditioned medium, L-929 cells were cultured in DMEM (Gibco) supplemented with 10% FBS (Sigma) and 1% penicillin-streptomycin (Gibco) for 10-14 days. The culture supernatant was collected, centrifuged to remove cells and debris, and further filtered through a 0.22 μm PVDF filter (Corning). Bone marrow cells were then cultured in DMEM (Gibco) supplemented with 10% FBS, 1% penicillin-streptomycin, and 25% L-929 conditioned medium in ultra-low attachment dishes at 37 °C in a humidified incubator with 5% CO₂. To induce differentiation into BMDMs, the medium was refreshed on day 3, and cells were cultured for a total of 5 days. Differentiated BMDMs were detached using 1 mM EDTA and collected for subsequent experiments.

### 4.5 Reverse Transcription-Quantitative Polymerase Chain Reaction (RT-qPCR)

Differentiated young and old BMDMs were seeded in 6-well plates and cultured in a humidified incubator at 37 °C with 5% CO₂ for 3 days. Cells were then harvested, and total RNA was extracted using the Direct-zol™ RNA Miniprep Kit (Zymo Research, R2052) according to the manufacturer’s instructions. RNA concentration and purity were assessed using a NanoDrop spectrophotometer (Thermo Scientific). Complementary DNA (cDNA) was synthesized from total RNA using the High-Capacity cDNA Reverse Transcription Kit (Applied Biosystems™, 4368814) following the manufacturer’s protocol. Quantitative polymerase chain reaction (qPCR) was subsequently performed with SYBR Green chemistry, and fluorescence signals were detected on the LightCycler 480 system using SYBR Green I Master Mix (Roche, 04887352001).

### 4.6 Mitochondrial Metabolism

To assess oxidative phosphorylation (OXPHOS) function, 5 × 10^4^ BMDMs were seeded in Seahorse XF 96-well microplates (Agilent) and cultured for 24 hours, and further prepared for Mito stress test analysis according to the manufacturer’s instructions for the Seahorse XF Cell Mito Stress Test Kit (Agilent, 103015-100). The oxygen consumption rate (OCR) was subsequently measured using a Seahorse XF Analyzer (Agilent).

### 4.7 Co-culture Experiments and Flow cytometry

For phagocytosis assays, cancer cells were labeled with Tag-it Violet™ Proliferation and Cell Tracking Dye (1:1000; BioLegend, 425101) for 20 min at 37 °C prior to co-culture with BMDMs or RAW 264.7 cells. The co-culture duration and cell ratios varied depending on the experimental design and are specified in the corresponding figure legends. For STAT6 inhibition, RAW 264.7 cells were pretreated with 2 μM AS1517499 (STAT6 inhibitor; Selleckchem, S8685) for 3 days, with medium refreshed daily, prior to co-culture with cancer cells. After co-culture, cells were harvested using either 1 mM EDTA (Sigma) or 0.25% trypsin (Gibco) and prepared for flow cytometry analysis.

For flow cytometry, single-cell suspensions were washed with PBS containing 2% FBS and stained with fluorophore-conjugated antibodies, including Zombie NIR™ Fixable Viability Dye (1:500; BioLegend, 423106) and anti-F4/80 (1:50; BioLegend, 111704), for 20 min on ice. Fluorescence signals were acquired using a BD FACSCelesta Cell Analyzer (BD Biosciences), and data were analyzed with FlowJo software (BD Biosciences). Macrophage-cancer cell fusion events were quantified as the percentage of Tag-it Violet+/F4/80+ cells within the Zombie NIR- (live-cell) population. Representative gating strategies are provided in Supplementary Figure S1.

### 4.8 Multiplex Immunofluorescence Analysis

For co-culture experiments, cells were seeded onto poly-L-lysine-coated slides and fixed with 4% paraformaldehyde. Following fixation, samples were washed with Multistaining Buffer (Lunaphore) and permeabilized with 0.1% Triton X-100 for 10 min. Cyclic immunofluorescence staining was performed using the Lunaphore COMET™ platform according to the manufacturer’s instructions. Signal intensities for each antibody were quantified using HORIZON™ software (Lunaphore) and further analyzed in RStudio (Posit Software, PBC). Primary antibodies used in this study included: SDHB (1:100, NBP1-87069; Novus Biologicals); Ki-67 (1:150, NBP2-80822; Novus Biologicals); ATP5a (1:100, ab14748; Abcam); ATP5d (1:100, PA5-21361; Invitrogen); and F4/80 (1:100, #70076; Cell Signaling Technology). Secondary antibodies used in this study included: Alexa Fluor 647-conjugated goat anti-rabbit (1:200, A32733, Thermo Fisher Scientific), Alexa Fluor 555-conjugated goat anti-rabbit (1:100, A32732, Thermo Fisher Scientific), Alexa Fluor 555-conjugated goat anti-mouse (1:100, A32727, Thermo Fisher Scientific), and Alexa Fluor 647-conjugated goat anti-rat (1:200, A48265, Thermo Fisher Scientific).

### 4.9 Statistical Analysis

Data were obtained from at least three independent experiments and are presented as mean ± standard error of the mean (SEM). Statistical comparisons between groups were performed using a two-tailed Mann-Whitney U test in GraphPad Prism (GraphPad Software). A *p*-value < 0.05 was considered statistically significant.

## Author Contributions

Conceptualization, T.H.M.H., and C.N.H. ; Methodology, L.Y.W., C.W.C., C.C.C., and H.C.L.; Software, L.Y.W., H.C.L., C.W.C., and C.N.H.; Validation, L.Y.W., H.C.L., and C.W.C.; Formal Analysis,., L.Y.W., H.C.L., and C.N.H.; Investigation, L.Y.W., H.C.L., and C.W.C.; Resources, T.H.M.H. and C.N.H.; Writing – Original Draft, L.Y.W., H.C.L., C.W.C., T.H.M.H., and C.N.H.; Writing - Review & Editing, all authors; Funding Acquisition and Supervision, T.H.M.H. and C.N.H.

## Acknowledgments

The authors acknowledge the Bioanalytics and Single-Cell Core for multiplex immunofluorescence imaging.

## Funding

This work was supported by Nathan Shock Center of Excellence in the Biology of Aging (C.N.H), NIH U01 CA283749 and pilot grant (T.H.M.H.), P30 CA054174, and the Max and Minnie Tomerlin Voelcker Fund (T.H.M.H.).

## Conflicts of Interest

The authors declare no conflicts of interest.

## Data Availability Statement

The data that support the findings of this study are available in the Supporting Information of this article.

## Notes

### Competing Interest Statement

The authors have declared no competing interest.

