## Supplemental information for "Aging-Associated Decline in Macrophage STAT6-OXPHOS Programs Promotes Tumor-Like Multinucleated Syncytia"

| Chia-Nung Hung[1]

**Table S1. Clinicopathological characteristics of patients with bone metastases included in this study.**

| Patient number | TNM staging of initial diagnosis | Age at bone metastasis* | Location of bone metastasis | Diagnosis |
|---|---|---|---|---|
| Breast cancer | | | | |
| B1 | T4b,N0,M1 | 47 | Left femur | Invasive ductal carcinoma |
| B2 | pT2 N0 M0 | 73 | Right proximal femur | Invasive ductal carcinoma |
| B3 | T2,N1a,M0 | 53 | N/A | Invasive ductal carcinoma |
| B4 | T2,N2a,M0 | 64 | Left femur | Invasive lobular carcinoma |
| B5 | T2,N1,M1 | 78 | Cervical spine | Invasive lobular carcinoma |
| B6 | T2,N0,M0 | 47 | Thoracic spine | Invasive ductal carcinoma |
| B7 | T1c,N2,M0 | 60 | Humerus | Mucinous adenocarcinoma |
| B8 | N/A | 68 | Cervical spine | Invasive ductal carcinoma |
| B9 | pT1c,N0,M0 | 57 | Cervical and thoracic spine | Invasive ductal carcinoma |
| B10 | T3 N0 M0 | 58 | Humerus | Invasive ductal carcinoma |
| Lung cancer | | | | |
| L1 | T2a,N0,M1c | 81 | Right femoral shaft | Adenocarcinoma |
| L2 | T2a,N1,M1c | 61 | Left humerus | Squamous cell carcinoma |
| L3 | T4,N3,M1c | 75 | Right femoral shaft | Adenocarcinoma |
| L4 | pT1aN0M0 | 69 | Right acromion | Adenocarcinoma |
| L5 | T4,N3,M1c | 58 | Right femur | Adenocarcinoma |
| L6 | T4,N2,M1c | 71 | Left femur | Adenocarcinoma |
| L7 | T2a,N0,M1b | 63 | Humerus | Non-small cell carcinoma |
| L8 | T2a,N3,M1b | 48 | Femoral bone, left | Non-small cell carcinoma |
| L9 | Tx,N3,M1a | 57 | Lumbar spine | Adenocarcinoma |
| L10 | pT3,N2,M1a | 63 | Pelvis bone, right | Adenocarcinoma |
| L11 | Tx,Nx,M1b | 72 | Left pelvic | Adenocarcinoma |
| L12 | Tx,Nx,M1b | 77 | Left femur | Adenocarcinoma |
| L13 | T2a,N3,M1c | 61 | Lumbar spine | Adenocarcinoma |
| L14 | T4,N0,M1c | 59 | Thoracic spine | Adenocarcinoma |
| L15 | T4,N3,M1c | 62 | Thoracic spine | Squamous cell carcinoma |
| Prostate cancer | | | | |
| P1 | c:Tx Nx M1,IVA | 85 | Spine T4-5 | Metastatic carcinoma, compatible with metastatic prostatic carcinoma |
| P2 | c*:T3N1M1b,IV | 85 | Spine L4 | Metastatic carcinoma, in favor of metastatic prostatic adenocarcinoma |
| P3 | cT3aN0M0 | 78 | Spine C5 | Metastatic carcinoma |
| P4 | c:Tx Nx M1,IV | 86 | Spine T5-6 | Metastatic carcinoma |
| P5 | c:T3a-bNxM1b,IVB | 92 | Spine S1 | Metastatic adenocarcinoma, compatible with prostatic origin |
| P6 | c:TxN1M1c,IVB | 57 | Spine T6-9 | Metastatic adenocarcinoma, consistent with prostatic origin |
| P7 | p:T4N1M1,IVB | 63 | Lumbar spine L3-5 | Metastatic carcinoma, compatible with prostate origin |

* Age of initial diagnosis is not provided.

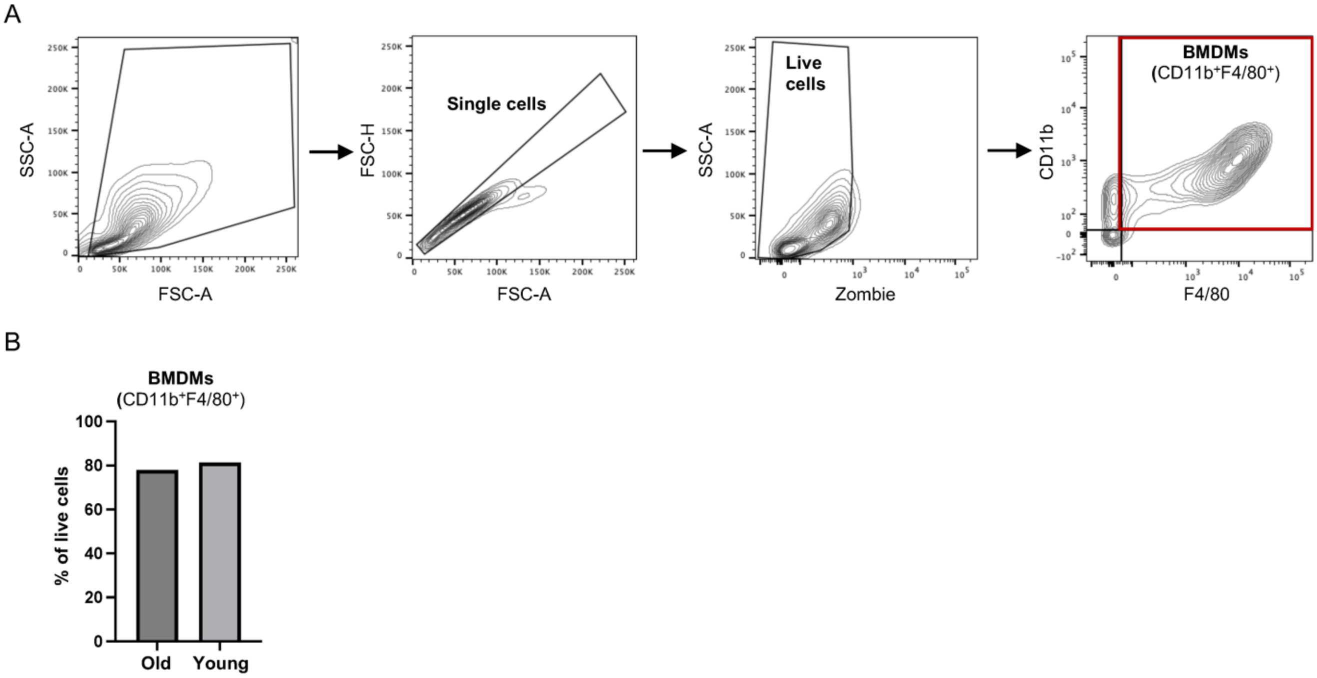


**FIGURE S1 | Flow cytometric characterization of BMDMs isolated from young and aged mice.** (A) Representative flow cytometry gating strategy used to identify bone marrow–derived macrophages (BMDMs). Cells were sequentially gated based on FSC-A/SSC-A, singlets (FSC-H versus FSC-A), live cells by Zombie dye exclusion, and CD11b^+/F4/80^+ macrophages. (B) Quantification of the percentage of CD11b^+/F4/80^+ BMDMs among live cells isolated from young and aged mice. Data are presented as the percentage of live cells.

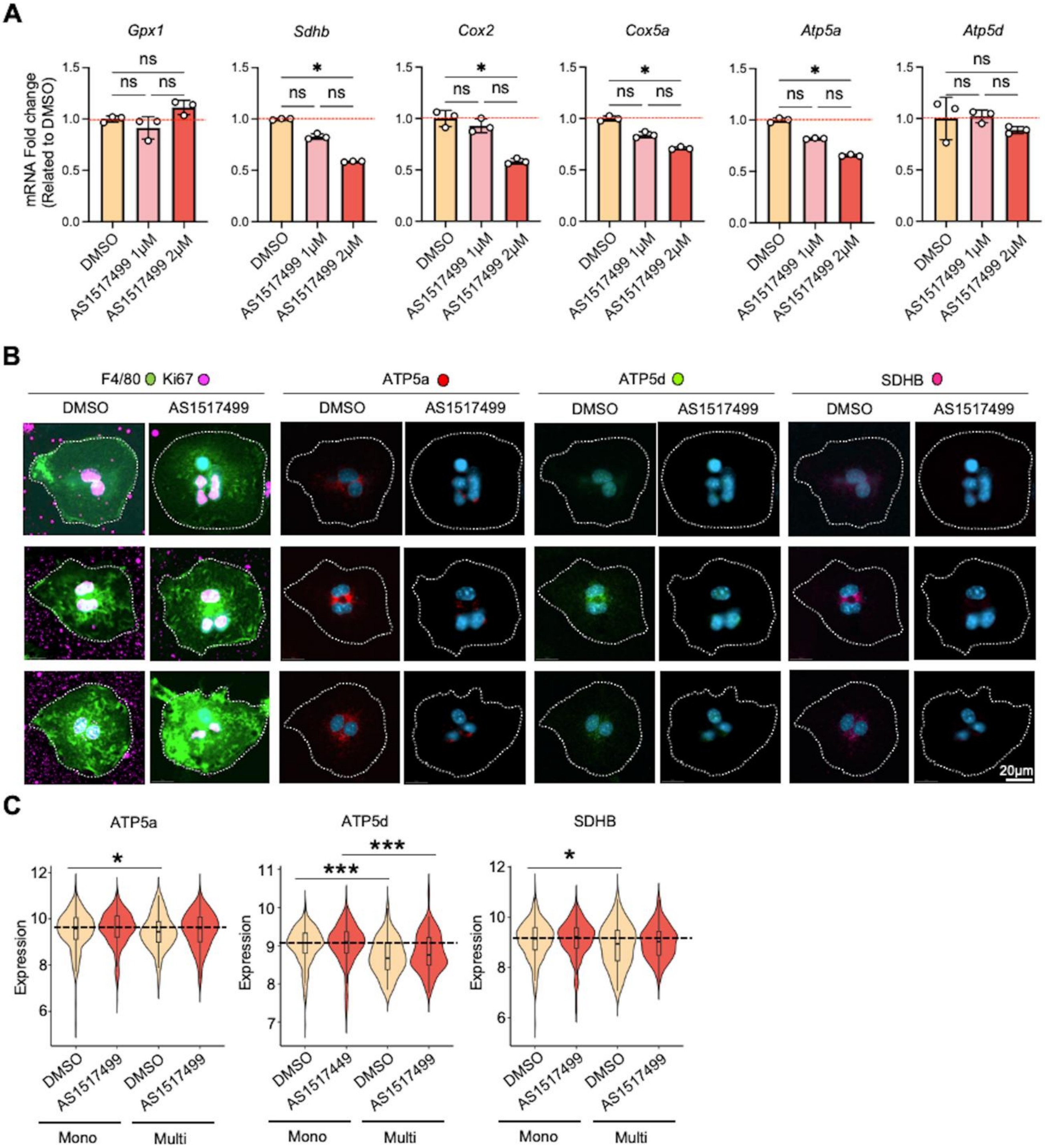

A
Gpx1
Sdhb
Cox2
Cox5a
Atp5a
Atp5d
mRNA Fold change (Related to DMSO)
ns
*
DMSO
AS1517499 1μM
AS1517499 2μM
B
F4/80 Ki67
ATP5a
ATP5d
SDHB
DMSO
AS1517499
20μm
C
ATP5a
ATP5d
SDHB
Expression
*
***
DMSO
AS1517499
AS1517449
Mono
Multi

**FIGURE S2 | Effects of STAT6 inhibition on OXPHOS-associated gene and protein expression in macrophage–tumor fusion cells.**

(A) Quantitative PCR analysis of OXPHOS-associated genes, including *Gpx1, Sdhb, Cox2, Cox5a, Atp5a,* and *Atp5d*, in macrophage–tumor co-cultures treated with DMSO, AS1517499 (1 μM), or AS1517499 (2 μM). Gene expression levels were normalized to the DMSO group. Data are presented as mean ± SEM (n = 3 per group). Statistical significance was determined by ordinary one-way ANOVA. $*P < 0.05$; *ns, not significant.* (B) Representative multiplex immunofluorescence images of macrophage–tumor fusion cells treated with DMSO or AS1517499 and stained for F4/80, Ki67, ATP5a, ATP5d, and SDHB. Dashed lines indicate cell boundaries. Scale bar, 20 μm. (C) Quantification of ATP5a, ATP5d, and SDHB expression in mononucleated (Mono) and multinucleated (Multi) fusion cells treated with DMSO or AS1517499. Data are presented as violin plots showing distribution and median values. Statistical significance was determined by the Mann-Whitney test. $*P < 0.05$; $***P < 0.001$.

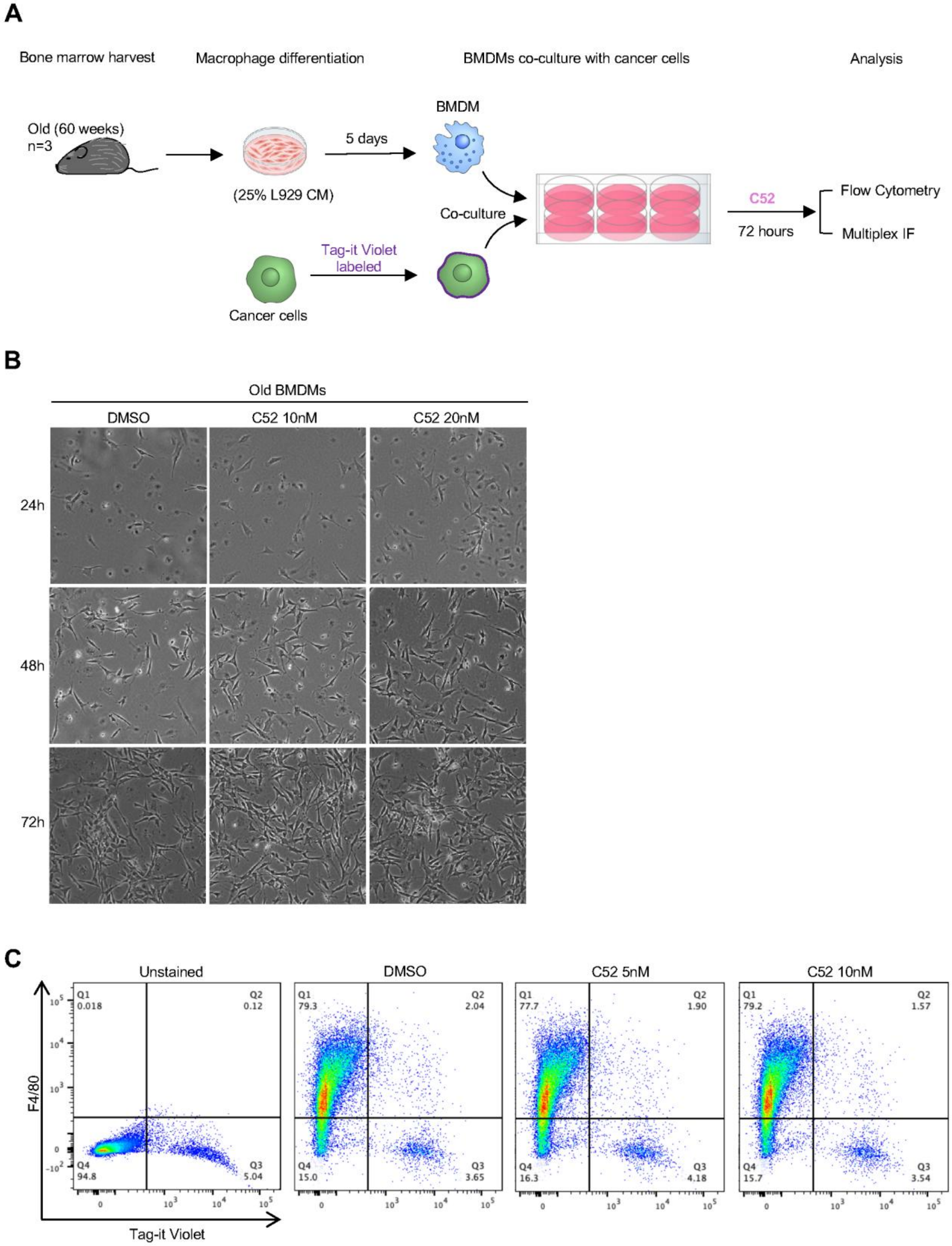
A
Bone marrow harvest
Macrophage differentiation
BMDMs co-culture with cancer cells
Analysis
Old (60 weeks)
n=3
5 days
BMDM
(25% L929 CM)
Co-culture
C52
72 hours
Flow Cytometry
Multiplex IF
Tag-it Violet
labeled
Cancer cells
B
Old BMDMs
DMSO
C52 10nM
C52 20nM
24h
48h
72h
C
Unstained
DMSO
C52 5nM
C52 10nM
F4/80
Tag-it Violet
Q1
0.018
Q2
0.12
Q4
94.8
Q3
5.04
Q1
79.3
Q2
2.04
Q4
15.0
Q3
3.65
Q1
77.7
Q2
1.90
Q4
16.3
Q3
4.18
Q1
79.2
Q2
1.57
Q4
15.7
Q3
3.54

**FIGURE S1 | Experimental workflow and characterization of C52-treated aged BMDM–tumor cell co-cultures.**

(A) Schematic illustration of the experimental design. Bone marrow cells isolated from aged mice (60 weeks old, n = 3) were differentiated into bone marrow–derived macrophages (BMDMs) in medium containing 25% L929 conditioned medium for 5 days. BMDMs were subsequently co-cultured with Tag-it Violet-labeled cancer cells in the presence of the EP2 antagonist C52 for 72 h prior to flow cytometric and multiplex immunofluorescence analyses. (B) Representative bright-field images of aged BMDM co-cultures treated with DMSO, C52 (10 nM), or C52 (20 nM) at 24, 48, and 72 h after co-culture. (C) Representative flow cytometry plots showing F4/80 and Tag-it Violet staining in co-cultures treated with DMSO, C52 (5 nM), or C52 (10 nM). Unstained cells were used as controls for gating.
